# Ultra-fast genetic colocalisation across millions of traits

**DOI:** 10.1101/2025.08.25.672103

**Authors:** Mihkel Jesse, Ago-Erik Riet, Kaur Alasoo

**Affiliations:** Institute of Computer Science, University of Tartu, Tartu, Estonia; Institute of Mathematics and Statistics, University of Tartu, Tartu, Estonia

## Abstract

Colocalisation is a powerful approach to assess if two genetic association signals are likely to share a causal variant. However, association analyses in large biobanks and molecular quantitative trait loci (molQTL) studies now routinely identify millions of association signals across thousands of traits, making it infeasible to test for colocalisation between all pairs of signals. Here we introduce *gpu-coloc*, a GPU-accelerated re-implementation of the coloc algorithm that combines efficient data storage with parallelisation to achieve a 1000-fold speed increase while maintaining near-identical results. As a result, the run time of gpu-coloc now approaches the colocalisation posterior probability (CLPP) method, a competing method that only uses information from fine mapped credible sets to detect colocalisations. Using summary statistics from UK Biobank, FinnGen, and eQTL Catalogue, we demonstrate that gpu-coloc and CLPP detect highly concordant results, especially when restricting the analysis to confidently fine mapped signals. We introduce the colocalisation collider metric to quantify spurious colocalisations in large-scale colocalisation graphs and use it to choose decision thresholds that provide a reasonable trade-off between sensitivity and specificity. Finally, we demonstrate how gpu-coloc can also be applied to marginal GWAS summary statistics from studies that lack fine mapping, where it is still able to recover molQTL colocalisations for ∼80% of the GWAS loci. Our efficient software and comprehensive analyses provide practical guidelines for future large-scale colocalisation analyses.

## Introduction

Genome-wide association studies (GWAS) have identified millions of associations linking genetic variants to thousands of human traits and diseases. However, over 90% of these variants are non-coding and can often have cell type and context-specific effects, complicating efforts to identify their functional roles.^1,2^ Genetic colocalisation methods can help to interpret GWAS by identifying molecular traits and biomarkers that share causal variants with disease or trait GWAS signals.^3–7^ For example, colocalisation with gene expression quantitative trait loci (QTLs) has helped to prioritise effector genes and relevant cell types for various diseases.^8,9^ Thus, large-scale genetic colocalisation has the potential to greatly improve GWAS interpretation.

One of the most widely used colocalisation methods is coloc^3^ that only requires marginal association summary statistics from both studies, but needs to make a restrictive assumption of at most one causal variant per locus. With statistical fine mapping, it is possible to distinguish between multiple conditionally distinct signals at the same locus.^10,11^ These distinct signals are then summarised as credible sets, which are defined as the minimal set of variants that are expected to contain the (conditionally distinct) causal variant with certain probability (e.g. 95%). The colocalisation posterior probability (CLPP) method first presented in the eCAVIAR paper performs colocalisation at the level of credible sets, thus detecting colocalisation between individual fine mapped signals.^4^ However, the exhaustive search fine mapping algorithm used by eCAVIAR made it too slow for most practical datasets.^4,10^ Large-scale fine mapping became more affordable with the development of the highly scalable FINEMAP^12^ and Sum of Single Effects (SuSiE)^13^ algorithms. Importantly, both the coloc and CLPP methods have been adopted to support SuSiE output,^6,14^ making it possible to use fine mapped credible sets and signal-specific Bayes factors (BFs) from SuSiE directly in colocalisation.

Although a large proportion of GWAS summary statistics in the GWAS Catalog now follow a standard format,^15,16^ sharing of fine mapping results is much more fragmented. For example, Million Veterans Program^17^ and Open Targets Platform^9,18^ have only released posterior inclusion probabilities (PIPs) of the credible set variants that can be used by the CLPP colocalisation method. In contrast FinnGen^19^, eQTL Catalogue^20,21^ and a few other studies^14,22^ have published the logarithms of Bayes factors (LBFs) for all tested variants required by coloc^6^. Moreover, while CLPP is defined at the variant level, posterior probability of colocalisation (PP.H4) in coloc is defined at the locus level, making it tricky to compare results from these two methods to each other. Thus, rigorous empirical benchmarking is needed to understand the relative performance of the CLPP and coloc methods in identifying colocalising signals, and whether the ∼1000-fold additional space required to store signal-specific LBFs is justified by potentially increased sensitivity of coloc.

A key challenge in benchmarking colocalisation is computational efficiency. While CLPP can be calculated instantaneously, and has been used previously to perform biobank-scale colocalisations,^14,19^ the current R implementation of coloc does not scale well to millions of colocalisation tests. Under single causal variant assumption, coloc first converts marginal summary statistics to approximate Bayes factors (ABFs).^3^ In presence of multiple causal variants, this step is replaced by fine mapping with SuSiE to obtain signal-specific conditionally distinct LBFs. In the next step, both implementations (hereinafter coloc.abf and coloc.susie) then use the same algorithm (coloc.bf_bf, see Methods) to calculate coloc posterior probabilities between the two signals of interest. While the ABFs and LBFs can be pre-computed once and cached for all future colocalisation tests, the calculation of posterior probabilities by coloc.bf_bf needs to be repeated for all pairs of traits and is prohibitively expensive when performing millions of tests.

We developed gpu-coloc, a re-implementation of the coloc algorithm that combines caching of pre-computed Bayes factors (either ABFs or LBFs) with ultra-fast parallel calculation of coloc posterior probabilities on GPUs to achieve approximately 1000-fold speed-up over the original R implementation while yielding nearly identical results. We then applied gpu-coloc to fine mapped signals from the eQTL Catalogue^20,21^, FinnGen^19^ and Rahu *et al*., 2024^22^ (Figure 1). The fast re-implementation allowed us to systematically compare the colocalisation results from CLPP and gpu-coloc to empirically identify thresholds at which CLPP and gpu-coloc yielded comparable results. We found that when restricting the analysis to confidently fine mapped signals, >90% the colocalisation identified by CLPP and gpu-coloc were shared, demonstrating good concordance between the two methods. Finally, we demonstrate that gpu-coloc can also be used in the absence of fine mapping, in which case it is able to identify colocalising molecular quantitative trait loci (molQTLs) for ∼80% of the GWAS loci (compared to when full fine mapping results are available).

**Figure 1.**
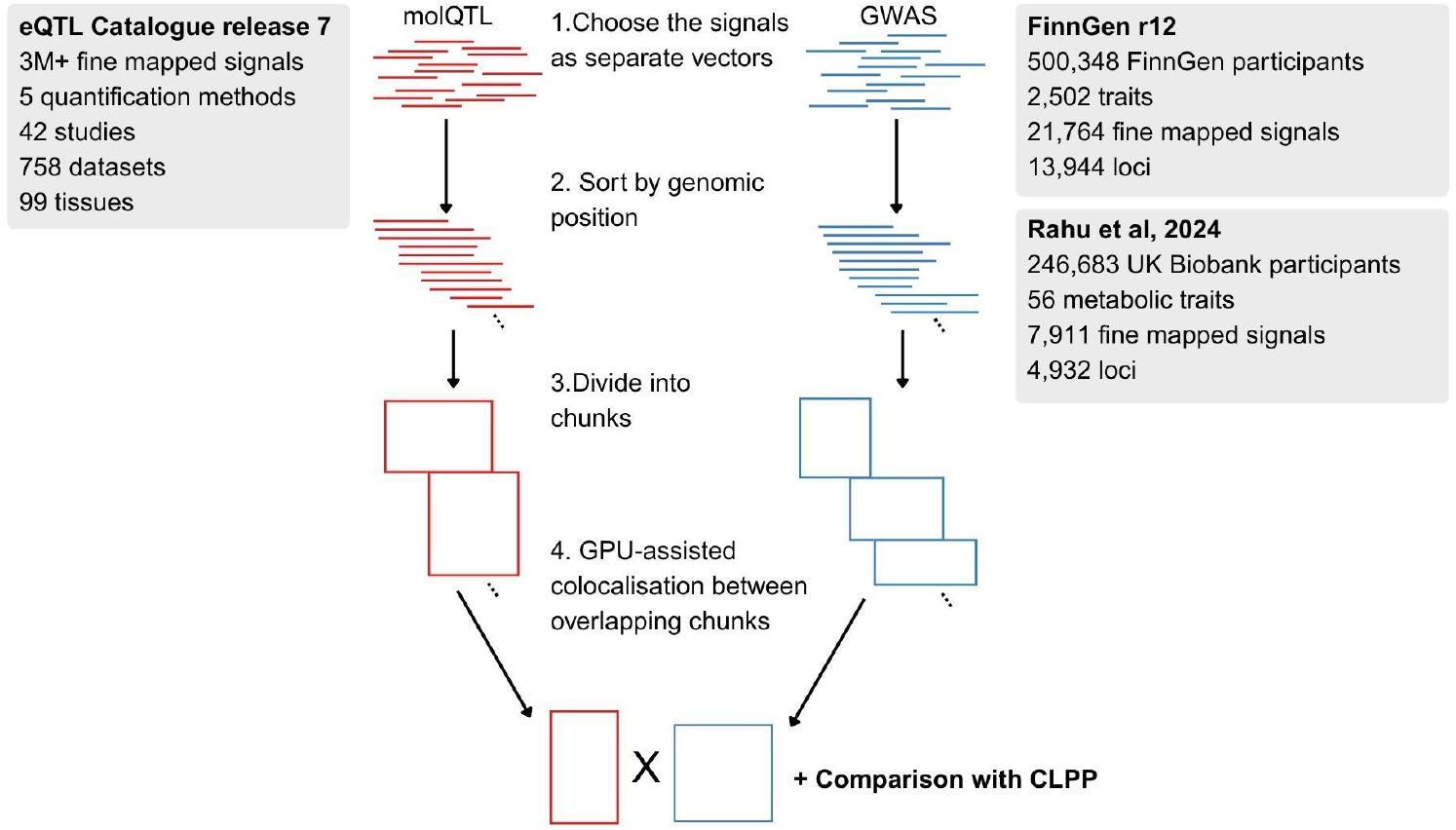
Overview of the gpu-coloc algorithm and our benchmark study. gpu-coloc extracts all independent associations from molecular QTL (molQTL) compendia or GWAS databases (1), sorts the signals by genomic position (2), divides the signals into chunks of ∼1000 signals (3), and performs GPU-assisted colocalisation between all overlapping chunks (4). We apply gpu-coloc to fine mapped association signals from eQTL Catalogue^20,21^, FinnGen^19^ and Rahu *et al*., 2024^22^, and compare the results to the colocalisation posterior probability (CLPP) approach.

## Results

### Implementation of the gpu-coloc method

Our method, gpu-coloc, is an efficient reimplementation of the coloc.bf_bf^3,6^ algorithm in Python and with GPU support. Instead of performing fine mapping at each GWAS locus, gpu-coloc uses SuSiE fine mapped LBFs from publicly available sources such as eQTL Catalogue^20,21^, FinnGen^19^ and Rahu *et al*., 2024^22^. These LBF vectors are first converted into matrices in parquet format for fast retrieval (Figure 1, see Methods). Alternatively, gpu-coloc can also leverage pre-calculated ABFs from studies that lack reliable fine mapping information. Similarly to coloc, gpu-coloc then estimates the posterior probability of whether two signals share a causal variant in the region (PP.H4, see Methods). Whereas the original coloc.bf_bf implementation in R tests for colocalisation between one pair of signals at a time, gpu-coloc can test thousands of pairs concurrently due to GPU-enabled matrix calculation (Figure 1). Massive matrix calculation is made possible by chunking signals by genomic position, where the matrix rows are signals and columns are variants, with the matrix entries being LBFs or ABFs. Our gpu-coloc implementation uses the same prior probability parameters as coloc.bf_bf: the priors *p*_1_ and *p*_2_ specifying the prior probabilities that any variant in the shared region is causal for the first or second signal, respectively; and the joint prior *p*_12_ specifying the prior probability that a variant is causal for both signals.^3^

An important consideration in any colocalisation analysis is how to handle associations with variants that are present in one dataset but missing in the other. By default, the pairwise comparisons performed by coloc.abf and coloc.susie use the intersection of variants present in both studies. This ensures that information from missing variants is appropriately treated as missing, rather than evidence for or against colocalisation. However, this is not feasible when calculating coloc posterior probabilities as matrix operations in gpu-coloc (Figure 1). As a workaround, within each collection of association summary statistics (e.g. different traits within FinnGen), gpu-coloc replaces all missing LBF values with a very small value (−1×10^6^), effectively assuming that these variants are not associated with the trait of interest (see Methods). Next, when performing colocalisations between two collections of summary statistics or fine mapping results (e.g. eQTL Catalogue vs FinnGen), gpu-coloc, similarly to coloc, only keeps the intersection of variants that are present in both collections. This approach achieves high computational efficiency while ensuring that variants systematically missing between two different collections of summary statistics (e.g. due to differences in allele frequency thresholds or genotype imputation panels) are still appropriately treated as missing.

We believe that our imputation approach is well suited for collections of summary statistics where the proportion of missing variants is low, such as in biobank-wide GWAS studies^17,19,22,23^, where the same set of variants is tested against all traits. Similarly, in the eQTL Catalogue, the genotype data from all studies have been imputed against the same 1000 Genomes reference panel^24^, thus reducing the number of missing variants to a minimum. However, our imputation strategy might not be appropriate when assembling an heterogeneous set of summary statistics from e.g. GWAS Catalog^16^ into a single large collection.

### gpu-coloc is a faster version of coloc.bf_bf with comparable accuracy

First, we validated gpu-coloc by comparing it to the official R implementation of coloc.^6^ We compared gene expression QTLs (eQTLs) from eQTL Catalogue GTEx^25^ datasets and fine mapped GWAS signals for 56 metabolic traits from the UK Biobank analysed by Rahu *et al*., 2024.^22^ To speed up the analysis, we only retained signals from chromosome 20. Using conservative prior probabilities (*p*_*1*_*= p*_*2*_ *=* 1×10^−4^, *p*_*12*_ *=* 1×10^−6^), we found that both gpu-coloc and coloc.bf_bf produced almost identical PP.H4 values when focussing on high confidence colocalisation (PP.H4 ≥ 0.8) (standard deviation of 2.3×10^−5^, maximum absolute error 1.3×10^−4^, mean absolute error = 1.1×10^−5^) (Figure 2A). Across the full range of PP.H4 values, most values were still highly concordant (mean absolute error = 7.0×10^−5^), but there were nine outliers (0.018% of all tested pairs) where the absolute error exceeded 0.02 (Figure 2A). However, the gpu-coloc PP.H4 value was more conservative than coloc.bf_bf PP.H4 value, indicating that our approach of replacing missing LBFs with very small values is unlikely to produce false positive results.

**Figure 2.**
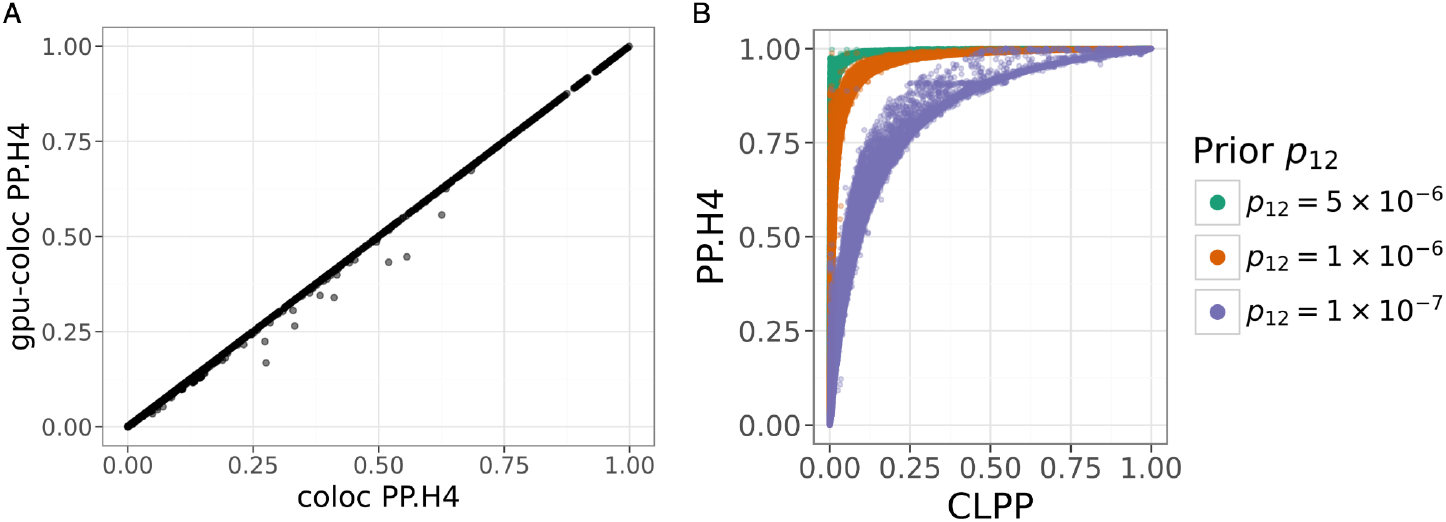
Comparison of the gpu-coloc, coloc.bf_bf, and CLPP colocalisation methods. (A) Comparison of the PP.H4 values between gpu-coloc and coloc.bf_bf. (**B**) Posterior probability comparison between gpu-coloc and CLPP across varying prior parameterisations (*p*_*12*_ = 5×10^−6^, 1×10^−6^, and 1×10^−^7).

In our benchmarks, gpu-coloc achieved approximately a 1000-fold speed-up compared to the standard R implementation. On the University of Tartu high performance computing (HPC) system, our implementation completed in 3 minutes and 20 seconds, approximately 887 times faster than the standard R version, which required 2 days, 1 hour, 19 minutes, and 17 seconds. On a M2 Max 14” MacBook Pro with 32 GB RAM, the performance difference was further amplified to a factor of 1150 (67 seconds versus 21 hours, 29 minutes, and 59 seconds). Formatting data for gpu-coloc, performed once prior to computation, took an additional 3 hours, 20 minutes, and 21 seconds on the HPC system, reducing the total effective runtime advantage to approximately 14.5-fold. However, the formatting step is computationally inexpensive, linear-time, and reusable across multiple analyses, thus imposing minimal overhead on subsequent studies. Furthermore, we have publicly released properly formatted LBF files for all eQTL Catalogue signals (250GB total, ∼50 GB per molQTL type) and will keep these files up-to-date as new versions of the eQTL Catalogue are released (see Data availability).

For testing large-scale colocalisation on a personal computer, we ran gpu-coloc between 306 eQTL Catalogue gene expression datasets (∼65 GB) and FinnGen r12 (∼13 GB) on a M2 Max 14” MacBook Pro with 32 GB RAM. The run took 1 hour, 9 minutes and 37 seconds, during which we tested colocalisation between 46,377,119 unique signal pairs. Thus, gpu-coloc enables researchers to test for colocalisation between their GWAS results and all eQTL Catalogue signals on their personal computers.

### Comparison of posterior probabilities between gpu-coloc and CLPP

An even faster alternative to coloc is the CLPP method.^4^ On the same benchmark where we compared gpu-coloc and coloc.bf_bf, gpu-coloc was ∼400 times slower (67 seconds) than CLPP (0.18 seconds). Furthermore, storing the LBFs for all tested variants and fine mapped signals required ∼1000-fold more space (1.6 GB) than simply storing the PIPs for the credible set variants (20.3 MB). However, previous studies have found it challenging to directly compare the two methods, because CLPP is defined at the variant level while PP.H4 is defined at the locus level.^26,27^ To explore this question in more detail, we leveraged our efficient gpu-coloc implementation to calculate both the CLPP and PP.H4 values for approximately 200 million molQTL and GWAS signal pairs across the eQTL Catalogue and FinnGen datasets. We found that although the relationship between the PP.H4 and CLPP values was not linear, there was a continuous lower bound allowing us to identify CLPP values above which the PP.H4 always exceeded a certain threshold (Figure 2B). For example, using coloc.bf_bf default prior probabilities of *p*_*1*_ = *p*_*2*_ = 1×10^-4^ and *p*_*12*_ = 5×10^−6^, and filtering for pairs where CLPP ≥ 0.01 revealed that all these signal pairs had PP.H4 ≥ 0.8 (Figure 2B). The converse was not always true as some signal pairs with PP.H4 ≥ 0.8 had lower (or missing CLPP) values (Figure 2B). Decreasing the *p*_*12*_ prior probability to 1×10^−6^ increased the CLPP threshold corresponding to PP.H4 ≥ 0.8 to 0.04. Finally, when the *p*_*12*_ prior probability was set to 1×10^−7^ then the CLPP value corresponding to PP.H4 ≥ 0.8 was ∼0.3 (Figure 2B). This demonstrates that it is possible to define thresholds where PP.H4 and CLPP are directly comparable, but leaves the question of choosing the optimal threshold and corresponding prior probabilities.

### Choosing the colocalisation thresholds for gpu-coloc and CLPP

If we decide to keep the PP.H4 threshold fixed (e.g. PP.H4 ≥ 0.8), then how should we select the optimal *p*_*12*_ prior probability and the corresponding CLPP threshold? For coloc, a recent analysis recommends setting *p*_*12*_ = 5×10^−6^ as a good starting point.^28^ Interestingly, at the PP.H4 ≥ 0.8 threshold, this corresponds to CLPP ≥ 0.01, which was independently shown in the CLPP method simulations to have a low false positive rate.^4^ However, these thresholds have been derived in the context of comparing a single GWAS locus to a single molQTL locus and it is unclear if they are still suitable when performing millions of colocalisations tests across hundreds of thousands of fine mapped loci. Furthermore, in cases of limited statistical power, SuSiE might sometimes merge the associations driven by two independent causal variants into a single signal. This is most obvious in eQTL studies where the same gene might have two or more conditionally distinct signals in a well-powered study but only one signal in a study with a smaller number of samples.^27^ This can then lead to *colocalisation colliders*, where two fine mapped signals for gene X in dataset A do not directly colocalise with each other (since they are conditionally distinct), but do colocalise with a non-finemapped signal for the same gene in dataset B (Figure 3A). The simplest collider events contain three signals, but they could also be part of longer chains. While these colocalisation colliders might technically not be false positives (there is still at least one shared causal variant in both studies), they complicate the interpretation of conditionally distinct signals which might perturb the target gene via different mechanisms. Thus, it is desirable to reduce colocalisation colliders as much as possible.

**Figure 3.**
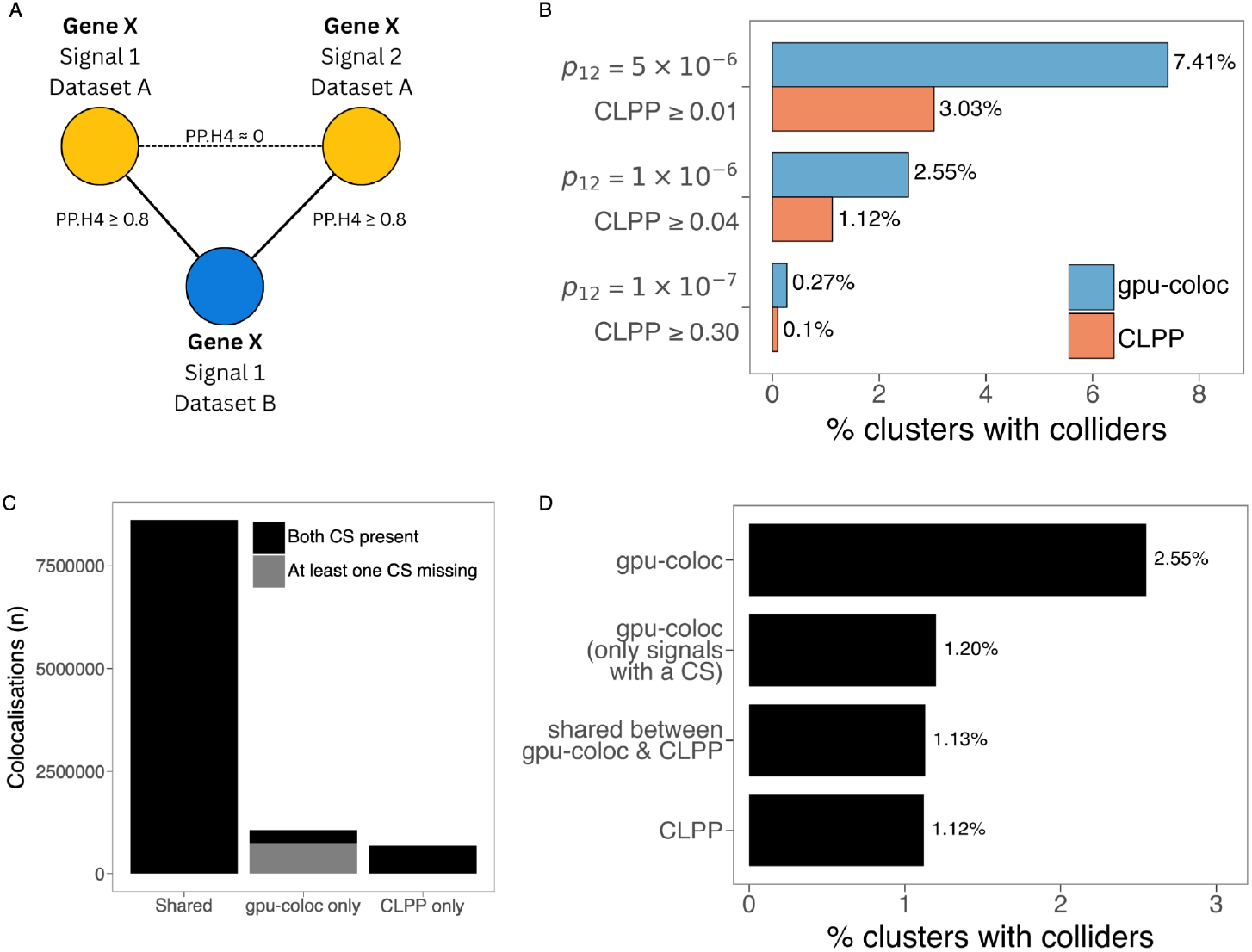
Colocalisation colliders and sources of differences. (**A**) Example of a colocalisation collider. Two fine mapped eQTL signals for gene X in dataset A (signals 1 and 2, top row) do not colocalise with each other (because they are conditionally distinct), but both separately colocalise with an eQTL signal for the same gene in dataset B. (**B**) Effect of *p*_*12*_ prior probability and CLPP threshold on the rate of colocalisation colliders observed in the eQTL Catalogue all-against-all eQTL colocalisation graph. For all *p*_*12*_ values, PP.H4 threshold was set to PP.H4 ≥ 0.8. (**C**) Number of method specific colocalisations results between gpu-coloc and CLPP inside the eQTL Catalogue all-against-all colocalisation at comparable decision thresholds (CLPP ≥ 0.04 and *p*_*12*_ = 1×10^−6^, PP.H4 ≥ 0.8), and the proportion of gpu-coloc specific results, where either signal did not have a credible set. (**D**) Percentage of clusters containing at least one colocalisation collider for all gpu-coloc colocalisation results (CLPP ≥ 0.04 and *p*_*12*_ = 1×10^−6^, PP.H4 ≥ 0.8), for gpu-coloc results where both signals have credible sets, for colocalisation results shared by gpu-coloc and CLPP, and all CLPP colocalisation results.

To identify colocalisation colliders, we used gpu-coloc and CLPP to perform all-against-all colocalisation between 636,057 fine mapped eQTL signals in the eQTL Catalogue^20,21^. We constructed a colocalisation graph, where the nodes are the signals and the edges are all pairwise significant colocalisation between the signals (PP.H4 ≥ 0.8). We then extracted all connected components (clusters) of that graph and counted the number of clusters that contained at least one colocalisation collider. At comparable thresholds, the rate of colliders was always higher for gpu-coloc relative to CLPP and decreased as the thresholds became more stringent (Figure 3B). For example, with *p*_*12*_ = 5×10^-6^ and PP.H4 ≥ 0.8, 7.41% of gpu-coloc clusters contained at least one colocalisation collider. In contrast, 3.03% of the clusters with CLPP ≥ 0.01 contained at least one collider. With *p*_*12*_ = 1×10^-6^ and corresponding CLPP ≥ 0.04, the rate of colliders decreased to 2.54% and 1.12%, respectively (Figure 3B). Since the rate of decrease in collider events was lower for more stringent CLPP values (Figure S1), we decided to use CLPP ≥ 0.04 (and corresponding *p*_*12*_ = 1×10^-6^) as our default thresholds for the remaining analyses.

### Colocalisations colliders are caused by fine mapping uncertainty

Next, we wanted to see what leads to the increased rate of colocalisation colliders in the gpu-coloc analysis compared to CLPP. In the eQTL Catalogue all-against-all colocalisation, we detected a total of 10,355,951 colocalisation events, 8,609,903 (83%) were shared between gpu-coloc and CLPP, 1,073,840 (10%) were detected only by gpu-coloc, and 672,208 (6%) were detected only by CLPP (Figure 3C). These proportions varied slightly in the eQTL Catalogue vs FinnGen and eQTL Catalogue vs Rahu *et al*., 2024 analyses (Figure S2). Notably, the majority (68%) of the additional colocalisation detected by gpu-coloc were between signals where at least one of them was missing a credible set (Figure 3C). The primary reason why SuSiE might not report a credible set in the presence of large LBFs (so that the colocalisation could be detected by gpu-coloc and not by CLPP) is when the variants to be included in the credible set would fail SuSiE’s purity filter. This filter is implemented in SuSiE to reduce the probability that a credible set would contain more than one independent causal variant. By default, this means that the minimal LD between any two variants included in the credible set has to be r^2^ > 0.5. Thus, we hypothesised that these SuSiE signals that failed purity filtering might have caused the increased rate of colocalisation colliders that we observed for gpu-coloc.

To test this, we excluded the 734,986 edges involving at least one SuSiE signal without a credible set (Figure 3C) from the gpu-coloc eQTL Catalogue all-against-all colocalisation graph and re-calculated the colocalisation collider rate. The collider rate decreased from 2.55% to 1.2% (Figure 3D). In contrast, when we excluded a random set of 734,986 edges 1000 times, the rate of clusters containing a collider decreased only marginally (new mean collider rate = 2.54%, p < 0.001). Finally, when we focussed on colocalisation that were shared between gpu-coloc and CLPP, then their collider rate was 1.13%, which was almost exactly the same as the collider rate of CLPP alone (1.12%) (Figure 3D). We observed broadly concordant results in the eQTL Catalogue vs FinnGen and eQTL Catalogue vs UK Biobank analyses (Figure S2). Thus, these results indicate that gpu-coloc colocalisations involving a SuSiE signal that failed purity filtering should be treated with caution. When all such signals were excluded from gpu-coloc then the proportion of shared colocalisation with CLPP increased from 83% to 89%, demonstrating that gpu-coloc and CLPP produce largely concordant results.

### Running gpu-coloc without fine mapping

By default, gpu-coloc expects SuSiE fine mapped signal-specific LBFs for each trait used in colocalisation. However, most publicly available GWASs (except for FinnGen and eQTL Catalogue) have not been fine mapped. Furthermore, accurate fine mapping might not be feasible for large-scale GWAS meta-analyses combining summary statistics from heterogeneous studies.^29^ In that scenario, it is possible to rely on the single causal variant assumption of the coloc.abf method^3^ and convert GWAS summary statistics into ABFs.

To test how the restrictive single causal variant assumption influences colocalisation, we obtained the marginal GWAS summary statistics from FinnGen r12^19^ and Rahu *et al*., 2024^22^, converted these to ABFs and then performed colocalisation against the fine mapped LBFs from the eQTL Catalogue (using the same prior *p*_*12*_ = 1×10^−6^). We then compared the ABF colocalisations to those found using the fine mapped LBFs available from both studies. We found that in both cases, fine mapping increased the number of colocalising GWAS loci by ∼18% and the number of conditionally distinct colocalising signals by >50% (Figure 4A). Nevertheless, for the majority of GWAS loci, fine mapping was not necessary to identify colocalisation with molQTLs, demonstrating that gpu-coloc can also be reliably used in scenarios where fine mapping results are not available.

**Figure 4.**
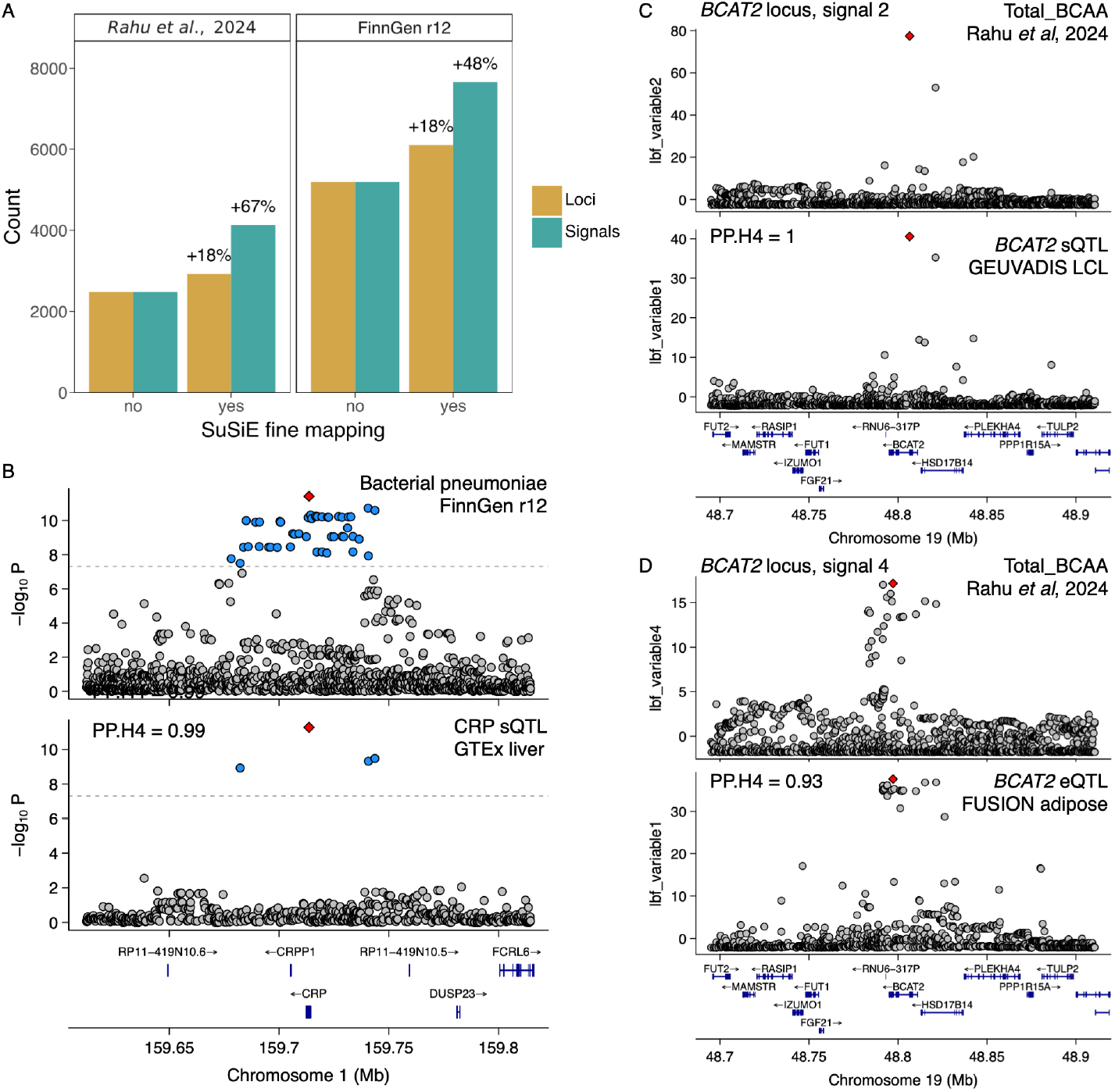
Fine mapping enables the detection of multiple independent colocalising signals at individual GWAS loci. (**A**) The number of GWAS loci and signals from Rahu *et al*, 2024 and FinnGen r12 that colocalise with at least one molecular QTL from the eQTL Catalogue (*p*_*12*_ = 1×10^−6^, PP.H4 ≥ 0.8). The results have been stratified by GWAS fine mapping status. (**B**) Example of a colocalisation detected without fine mapping. GWAS signal for bacterial pneumoniae at the *CRP* locus (1-159713648-C-G, rs1800947) colocalises with a *CRP* sQTL in the GTEx liver tissue dataset (QTD000270) from the eQTL Catalogue. (**C-D**) Examples of two colocalisation detected only after fine mapping. Secondary GWAS signal (**panel C**) for total branched-chain amino acids (Total_BCAA) (19-48806519-G-C, rs117048185) at the *BCAT2* locus colocalises with an sQTL for *BCAT2* in the GEUVADIS LeafCutter dataset (QTD000114) of the eQTL Catalogue. Fourth signal for Total_BCAA (19-48797174-G-A, rs35230038) at the *BCAT2* locus (**panel D**) colocalises with a *BCAT2* eQTL in the FUSION adipose tissue eQTL dataset (QTD000090) from the eQTL Catalogue.

For example, a GWAS hit for bacterial pneumoniae at the C reactive protein (*CRP*) locus (lead variant rs1800947) in FinnGen r12 colocalised with a splicing QTL for the *CRP* gene in the GTEx v8 liver dataset (QTD000270) (Figure 4B, Figure S3). As a result, the colocalisation was detected both with fine mapped LBFs (PP.H4 = 0.998) as well as with ABFs calculated from marginal summary statistics without fine mapping (PP.H4 = 0.988) (Figure 3B). Although *CRP* is a pattern recognition receptor that can bind to bacterial polysaccharides,^30^ it is also widely used as a diagnostic criterion to distinguish between bacterial and viral infections.^31,32^ Thus, it is highly likely that the FinnGen GWAS signal is better explained by diagnostic bias (i.e. if infected, variant carriers are more likely to receive diagnosis) rather than true biological effect on infection risk.

In contrast, at the branched-chain amino acid aminotransferase 2 (*BCAT2*) locus for total branched-chain amino acids (Total_BCAA) from Rahu *et al*., 2024^22^, there were five fine-mapped conditionally distinct signals that interfered with each other. Thus, using ABFs calculated from marginal summary statistics did not reveal any colocalisations. However, using fine mapped LBFs detected 22 colocalisations with four conditionally distinct signals (Table S1), including the two highlighted here: GWAS signal 2 (lead variant rs117048185) colocalised with a splicing QTL for *BCAT2* in the GEUVADIS^33^ lymphoblastoid cell line dataset (PP.H4 = 1) (Figure 4C, Figure S4) and GWAS signal 4 (lead variant rs35230038) colocalised with an eQTL for *BCAT2* in the FUSION^34^ adipose tissue dataset (PP.H4 = 0.93) (Figure 4D, Figure S5). Although rs117048185 is also annotated as a missense variant, it is located 2 bp from the splice donor site and is predicted by SpliceAI^35^ to lead to splice donor loss (delta score = 0.41), which is consistent with the reduced rate of exon 3 inclusion in the RNA-seq data (Figure S4). This highlights how fine mapping can reveal colocalisations with multiple conditionally distinct signals that perturb the same target gene via distinct mechanisms.

### Recommendations for large-scale colocalisation analysis

Based on our analysis, we make the following recommendations for future large-scale colocalisation analyses:

1. We believe that setting p_12_ prior to 1×10^−6^ and using PP.H4 ≥ 0.8 threshold for gpu-coloc represents a reasonable trade-off between sensitivity and limiting colocalisation colliders for most applications. For the CLPP method, this corresponds to retaining results with CLPP ≥ 0.04.
2. If PIPs from fine mapped credible sets are available for all traits of interest, then relying on the CLPP method alone is likely to be the most efficient approach with only minimal loss in sensitivity.
3. We recommend using gpu-coloc as a computationally efficient alternative to coloc.bf_bf when either some or all traits have not been fine mapped, or when there is a need to include low-confidence fine mapped signals that fail SuSiE’s purity filtering.

## Discussion

We introduce a novel implementation of the coloc algorithm, gpu-coloc, which enables biobank-scale colocalisation. Our approach achieves up to 1000-fold increase in computational speed compared to coloc.bf_bf while maintaining comparable accuracy. We performed colocalisation between all fine mapped molecular traits from eQTL Catalogue release 7, fine mapped GWAS signals from FinnGen release 12, and fine mapped metabolic traits from Rahu *et al*., 2024^22^. We identified thresholds at which gpu-coloc and CLPP yielded comparable results and introduced the concept of colocalisation colliders to quantify spurious colocalisations in large-scale colocalisation graphs. We found that although gpu-coloc had ∼20% increased sensitivity compared to CLPP, it also had ∼2.5-fold higher rate of colocalisation colliders. This difference was driven by low-confidence fine mapped signals that did not pass SuSiE’s purity filter and were thus excluded by CLPP. After these low-confidence fine mapped signals were excluded from gpu-coloc, both methods yielded highly concordant (∼90% shared) results. Finally, we demonstrate that gpu-coloc can still be used when fine mapping is not feasible for one (or both) sets of traits, in which case it is expected to find colocalisation for approximately 80% of the GWAS loci.

The gpu-coloc approach is related to a number of other methods. The tensorQTL^36^ software package also provides a GPU-enabled implementation of the coloc algorithm, but their implementation does not currently support SuSiE LBFs as input and is limited to testing colocalisation separately for each pair, thus missing a large proportion of the efficiency gains that we demonstrate. HyPrColoc^26^ is able to efficiently test the colocalisation between multiple signals at the same time, but is restricted by the single causal variant assumption and does not provide pairwise colocalisation estimates for all traits. Flanders only stores the LBFs for variants within the credible sets and imputes the others.^37^ Finally, gwas-pw^38^ and fastENLOC^5,7^ use enrichment analysis to identify prior probabilities for colocalisation instead of defining these values *a priori*.

Our approach also has several limitations. The GPU-enabled parallelisation of gpu-coloc assumes that there are multiple association signals in the same genetic locus. Thus, gpu-coloc performs best when testing for colocalisations between hundreds or even thousands of traits at the same time, or alternatively, when testing a single GWAS study against a large uniform collection of summary statistics. When testing for colocalisation between a single molQTL study and a single GWAS, then there is unlikely to be significant speed-up compared to the tensorQTL implementation. Furthermore, for one collection of summary statistics, gpu-coloc assumes that all variants were tested for all traits and that there are no missing values. This assumption is likely to be satisfied in the eQTL Catalogue and large, uniformly processed biobanks (e.g.

FinnGen^19^, Pan-UK Biobank^23^, Million Veterans Program^17^), but might be problematic when harmonising heterogeneous summary statistics from sources such as the GWAS Catalog^16^ or the IEU OpenGWAS database^39^.

We believe our comprehensive analysis provides practical guidelines for future large-scale colocalisation studies. We strongly encourage public sharing of fine mapping results whenever possible as this can help to distinguish between multiple conditionally distinct signals with different molecular mechanisms at the same locus (as illustrated by the *BCAT2* example in Figure 4). While sharing fine mapped LBFs for all tested variants does provide some advantages, our results indicate that ∼95% of high-confidence gpu-coloc colocalisations can already be identified with the CLPP method that requires only the PIPs from the credible set variants. Thus, we hope that studies that are currently unable to openly share full genome-wide summary statistics (e.g. due to re-identification risk), will be able to publicly release PIPs for the variants in fine mapped credible sets.

## Methods

### Datasets used for colocalisation analysis

The **eQTL Catalogue** is an open repository of uniformly processed human gene expression and splicing QTLs.^20,21^ Release 7 of the eQTL Catalogue contains 758 datasets from 42 studies, spanning 99 distinct tissues and/or cell types. The eQTL Catalogue contains over 3 million fine mapped signals, 2,142,311 of which have a credible set. The eQTL Catalogue is accessible at https://www.ebi.ac.uk/eqtl/.

The **UK Biobank** is a longitudinal biomedical study with approximately half a million voluntary participants aged 38-71 from the United Kingdom (at the time of recruitment between 2006 and 2010).^40^ In our colocalisation analyses we used GWAS summary statistics and fine mapping results from 246,683 UK Biobank analysed by Rahu *et al*. 2024.^22^ The study by Rahu *et al*. focused on 56 metabolomic biomarkers, which were measured from EDTA plasma samples between 2019 and 2024 using nuclear magnetic resonance platform from Nightingale Health. The fine mapped GWAS summary statistics contained 15,210 signals, from which 7,911 have credible sets.^22^ The dataset is available at https://doi.org/10.5281/zenodo.13821038.

The **FinnGen** release 12 is based on digitalized health data for 500,348 Finnish donors, mostly from hospital biobanks.^19^ The median age of the participant, while donating, was 53, with no age restrictions.The patients genotypes have been imputed using the Finnish population-specific SISu v.3 imputation reference panel.^19^ FinnGen release 12 has 13,944 genome-wide significant loci for 2,502 traits, 1,302 of which have been fine mapped. These loci contain 21,764 fine mapped signals, of which 17,844 have a credible set. The FinnGen dataset is accessible at https://www.finngen.fi/en/access_results.

### Implementation of gpu-coloc

gpu-coloc is a reimplementation of the coloc.bf_bf^6^ method in Python and with support for GPU acceleration using the Torch library. The original R implementation enables running coloc on two traits with multiple signals, but testing colocalisation is done consecutively on a CPU. We leverage the same idea, but instead of testing two signals of two traits at the same time with vector-based calculations, we use matrix-based calculations to leverage GPUs. Usually up to 10 signals are given for a locus. However, 10 x 10 tests are not yet effective, thus we chunk different signals in a region by their genomic position to create matrices with up to 1000 different signals. By default, colocalisation is then further divided into batches of 100 x 100 signals, but this can be further scaled with GPU memory (see below). We also implemented the trim_by_posterior function from coloc.bf_bf to omit results with low overlap of strongly associated SNPs. However it is important to keep in mind that Torch, and in extension gpu-coloc, supports only CUDA and Metal software for GPU calculation, thus gpu-coloc can be used on machines with an NVIDIA GPU (e.g. most HPCs, many Windows laptops and desktops) or M-series Macs.

### Data pre-formatting

GPU acceleration is enabled by pre-formatting signals into matrices. This first requires separating and summarising each signal (start and end of region, chromosome, highest LBF value). Here we discard all signals with maximum LBF < 5 (about ¾ of all signals), as they are unlikely to colocalise with other signals. Signals are sorted by the start of their region on every chromosome, then grouped into matrices based on overlap of regions. Since the signal regions are often defined around the lead variant or around a certain gene, then nearby signals overlap only partially. To format the regions into a matrix, we replace all missing LBF values with a placeholder value of −1×10^6^. This is a numerically negligible approximation of BF equals 0, thus assuming no association. Importantly, we assume that inside a collection of summary statistics (e.g. FinnGen or the eQTL Catalogue), all variants have been tested for all traits. Our approach is related to the idea of masking used to avoid confounding by multiple independent causal variants without fine mapping.^28^ Matrices are stored in parquet format, which provide both small file size and fast reading speed. Each matrix currently accommodates up to 1,000 signals.

### Calculating posterior probability of colocalisation (PP.H4)

Bayesian colocalisation methods such as coloc^3^ and gpu-coloc analyze a shared genetic locus consisting of n variants. Each trait under investigation is represented by an n-dimensional vector of LBFs (or ABFs), with each coordinate corresponding to a specific variant. For trait 1, we denote this vector by *a* = (*a*_1_, *a*_2_, …,*a*_*n*_), and for trait 2 by *b* = (*b*_1_, *b*_2_, …,*b*_*n*_). As Bayesian approaches, these methods require specifying prior probabilities: *p*_*1*_ for the probability that any given variant in the locus is associated with the first trait, *p*_*2*_ for the second trait, and *p*_*12*_ for association with both traits. To compute the posterior probability PP.H4, we first determine the unnormalised support for each hypothesis *H*0 through *H*4:

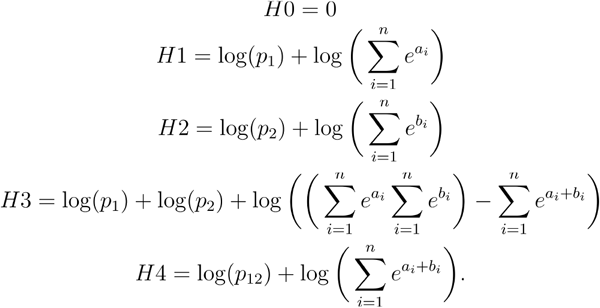

Once these supports are obtained, posterior probabilities can be calculated using:

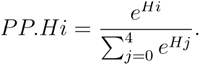

Most importantly

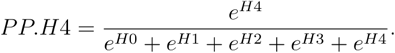

### Calculating colocalisation posterior probability (CLPP)

Similarly to coloc, CLPP^4^ assumes a shared locus consisting of n variants. For each trait, there is an n-dimensional vector where each coordinate corresponds to a variant and is assigned a PIP between 0 and 1. The PIP for a variant of a trait in a region can be approximated quite precisely by dividing its BF with the sum of all BFs for the trait in the region.^41^ Thus, while the LBFs used as input to coloc.bf_bf can be trivially converted to PIPs required by CLPP, CLPP as commonly implemented only uses PIPs for variants that are shared between two credible sets of interest. The sum of the PIPs for a vector equals 1. For trait 1, we denote this vector by *x* = (*x*_1_, *x*_2_, …,*x*_*n*_), and for trait 2 by *y* = (*y*_1_, *y*_2_, …,*y*_*n*_). CLPP is calculated as:

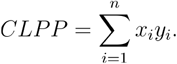

### Benchmarking

To benchmark the performance of gpu-coloc, we compared it with the original coloc.bf_bf implementation in R. Implementations were compared focusing on chromosome 20 between eQTL Catalogue gene expression datasets from GTEx^21^ and metabolic traits from Rahu *et al*., 2024^22^. We used prior probabilities *p*_*1*_ = *p*_*2*_ = 1×10^-4^ and p_12_ = 1×10^-6^. The R implementation was run using coloc on a CPU node in the University of Tartu HPC cluster, equipped with 1 AMD EPYC 7702 64-core processor, 60GB RAM, 8 TB SSD, and HDR Infiniband at 100 Gbps. gpu-coloc benchmark was conducted on a GPU node in the same HPC cluster, with an Intel Xeon E5-2650 v4 CPU, 60 GB of RAM, an NVIDIA Tesla V100 GPUs, with 16GB VRAM, and HDR Infiniband at 40 Gbps. We also tested both implementations on a MacBook Pro with an Apple M2 Max and 32 GB RAM.

### Colocalisations between eQTL Catalogue and FinnGen

We performed colocalisation between the whole of eQTL Catalogue release 7^21^ and all traits from FinnGen r12^19^. For the eQTL Catalogue, we used fine mapped LBFs available from the eQTL Catalogue FTP server. For FinnGen, we performed two separate analyses. First, we used fine mapped LBFs directly available from FinnGen. Secondly, to quantify the importance of fine mapping for detecting colocalisations, we also downloaded the marginal GWAS summary statistics from FinnGen and converted those to ABFs^42^ using the approx.bf.estimates (https://github.com/chr1swallace/coloc/blob/main/R/claudia.R#L96) function from the coloc R package.^3^ For colocalisation with fine mapped signals, we kept the priors *p*_*1*_ = *p*_*2*_ constant at 1×10^-4^ and varied *p*_*12*_ across *p*_*12*_ = 5×10^-6^, 1×10^-6^ and 1×10^-7^. For colocalisation with the non-finemapped approximate Bayes factors, we used the priors *p*_*1*_ = *p*_*2*_ = 1×10^-4^ and *p*_*12*_ = 1×10^-6^. For colocalisation with CLPP, we only included credible sets that passed SuSiE’s purity filter (low_purity == FALSE in FinnGen summary statistics). For comparison between the methods, we selected the CLPP thresholds as the lowest CLPP value rounded to two decimals, where PP.H4 for the selected prior was always at least 0.8.

### Colocalisations between eQTL Catalogue and metabolic traits

We performed colocalisation between the whole of eQTL Catalogue release 7^21^ and 56 metabolic traits from Rahu *et al*., 2024^22^. For gpu-coloc, we used priors *p*_*1*_ = *p*_*2*_ = 1×10^-4^ and *p*_*12*_ = 1×10^-6^. Fine mapping results released by Rahu *et al*. only contained credible sets that had already passed SuSiE’s purity filter.

### All-against-all colocalisation within the eQTL Catalogue

We used a subset of 306 eQTL Catalogue gene expression and protein QTL datasets, where the the quantification method in the dataset metadata table (https://github.com/eQTL-Catalogue/eQTL-Catalogue-resources/blob/master/data_tables/dataset_metadata.tsv) was either ‘ge’, ‘microarray’, or ‘aptamer’. These datasets contained a total of 2,748,413 conditionally distinct fine mapped signals, after discarding signals with maximum LBF < 5, 636,057 signals remained. We then tested for colocalisation between all pairs of signals, varying the *p*_*12*_ prior probability across *p*_*12*_ = 5×10^-6^, 1×10^-6^, and 1×10^-7^ while maintaining *p*_*1*_ = *p*_*2*_ = 1×10^-4^.

### Detecting colocalisation colliders

We define a colocalisation cluster as a connected component in the colocalisation graph built from colocalising signal pairs with PP.H4 ≥ 0.8. Nodes are fine mapped signals (dataset, trait, signal ID) and edges indicate a colocalisation (PP.H4 ≥ 0.8). A colocalisation collider is a cluster that contains at least two distinct signals of any single molecular trait (dataset, trait). For example, if signal 1 and signal 2 of trait A both colocalise with signal 2 of trait B, the set {A1, B2, A2} is a colocalisation collider (Figure 3A). The detection algorithm filters pairs to PP.H4 ≥ 0.8, builds the undirected graph on the remaining signals, finds connected components, and flags any component where any trait appears more than once. Signals that do not colocalise with any other signal are excluded from the analysis.

## Code availability

The source code of gpu-coloc is available at https://github.com/mjesse-github/gpu-coloc under MIT licence. The source code of coloc, which gpu-coloc derived from, is accessible at https://github.com/chr1swallace/coloc. Code used for writing this paper and generating the plots is accessible at https://github.com/AlasooLab/gpu-coloc-manuscript.

## Data availability

All data used for this article is available publicly: eQTL Catalogue release 7 at https://ftp.ebi.ac.uk/pub/databases/spot/eQTL/susie/, 56 metabolic traits by Rahu *et al*. 2024 at https://zenodo.org/records/13821038, and FinnGen release 12 at https://www.finngen.fi/en/access_results. Colocalisation results generated in this study are available at 10.5281/zenodo.15878809. The pre-formatted eQTL Catalogue input files for gpu-coloc are available from https://github.com/mjesse-github/gpu-coloc.

## Acknowledgements

We thank Ralf Tambets and Urmo Võsa for helpful comments on the manuscript. Some analyses presented in this paper were performed at the High Performance Computing Center, University of Tartu. M.J. and K.A. were supported by the Estonian Research Council (grant nos. PSG415 and MOB3ERC115). A-E.R. was supported by the Estonian Research Council (grant no PRG2531).

## Declaration of interests

The authors declare no competing interests.

## Supplementary Information

**Figure S1.**
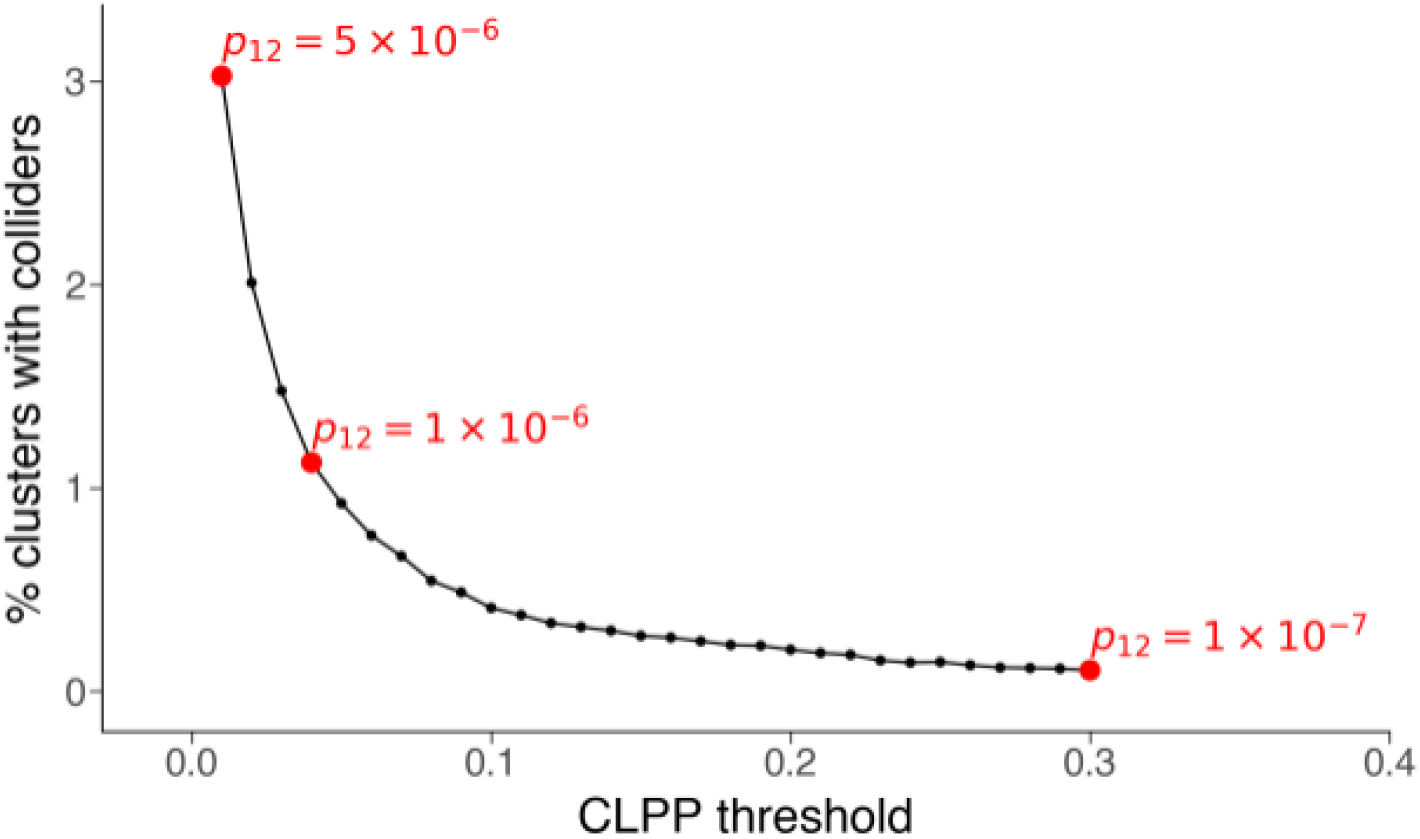
Effect of CLPP threshold on the rate of colocalisation colliders. The red dots illustrate the CLPP thresholds and *p*_*12*_ prior probabilities at which CLPP and gpu-coloc (PP.H4 > 0.8) produce comparable results.

**Figure S2.**
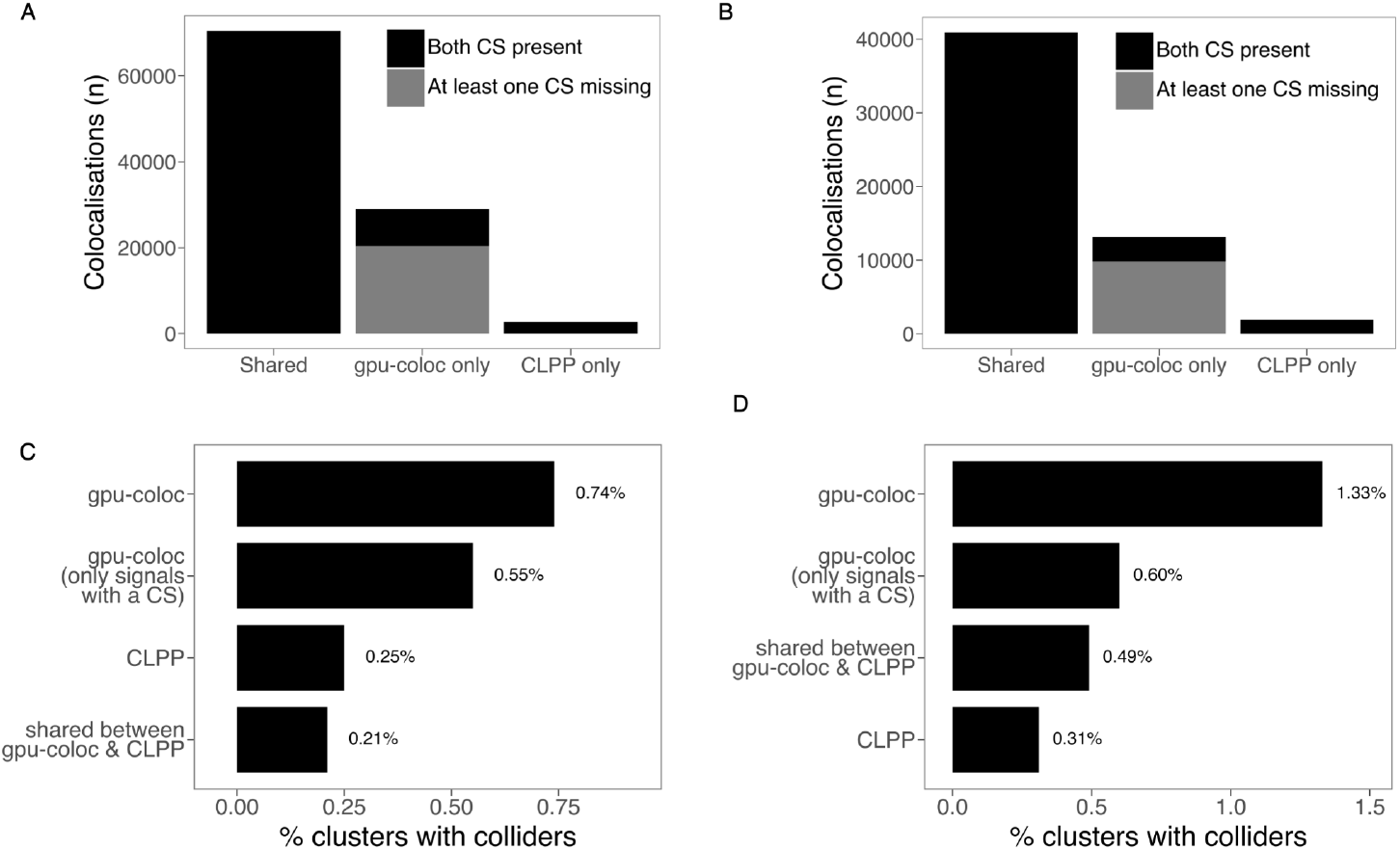
Sources of differences in colocalisation results between gpu-coloc and CLPP at comparable decision thresholds (CLPP≥0.04 and p_12_=1×10^−6^, PP.H4≥0.8). **(A)** Sources of difference from colocalisation between FinnGen *versus* eQTL Catalogue. **(B)** Sources of difference from colocalisation between 56 UK Biobank metabolic traits from Rahu *et al*., 2024 *versus* eQTL Catalogue. (**C-D**) Rate of colocalisation colliders in the eQTL Catalogue *vs* FinnGen r12 (**Panel C**) and eQTL Catalogue *vs* Rahu *et al*., 2024 (**Panel D**) analyses. Barplots show the percentage of clusters containing at least one colocalisation collider for all gpu-coloc colocalisation results (CLPP ≥ 0.04 and *p*_*12*_ = 1×10^−6^, PP.H4 ≥ 0.8), for gpu-coloc results where both signals have credible sets, for colocalisation results shared by gpu-coloc and CLPP, and all CLPP colocalisation results. CS - credible set.

**Figure S3.**
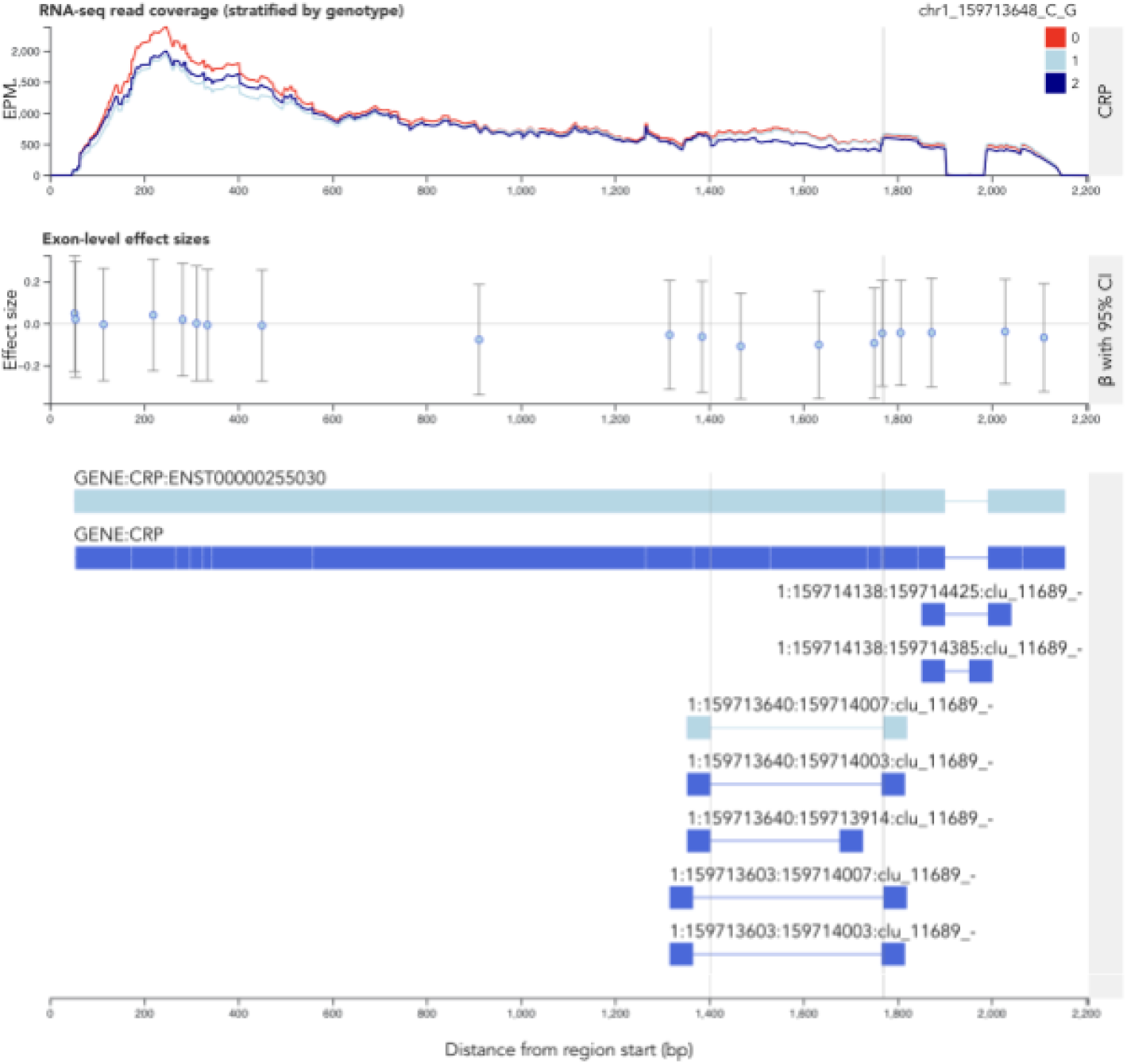
Visualisation of the *CRP* splicing QTL signal. RNA-seq read coverage across the *CRP* gene in the GTEx liver dataset (QTD000270) is stratified by the genotype of the lead sQTL variant (rs1800947). The interactive plot can be viewed in the ELIXIR-Estonia eQTL Catalogue Browser (<u>link</u>).

**Figure S4.**
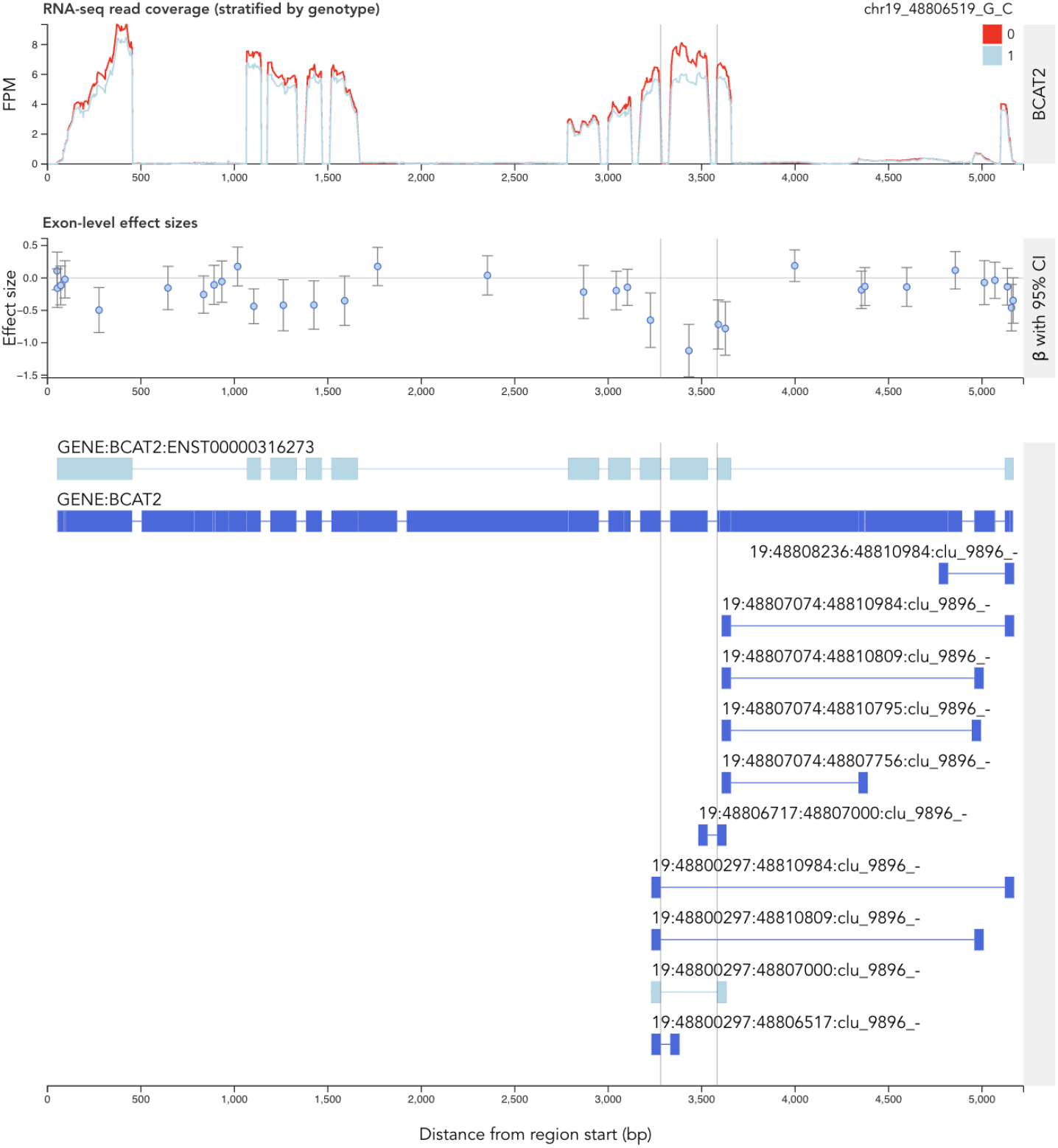
Visualisation of the *BCAT2* splicing QTL signal. RNA-seq read coverage across the *BCAT2* gene in the GEUVADIS lymphoblastoid cell line dataset (QTD000114) stratified by the genotype of the lead sQTL variant (rs117048185). The interactive plot can be viewed in the ELIXIR-Estonia eQTL Catalogue Browser (<u>link</u>).

**Figure S5.**
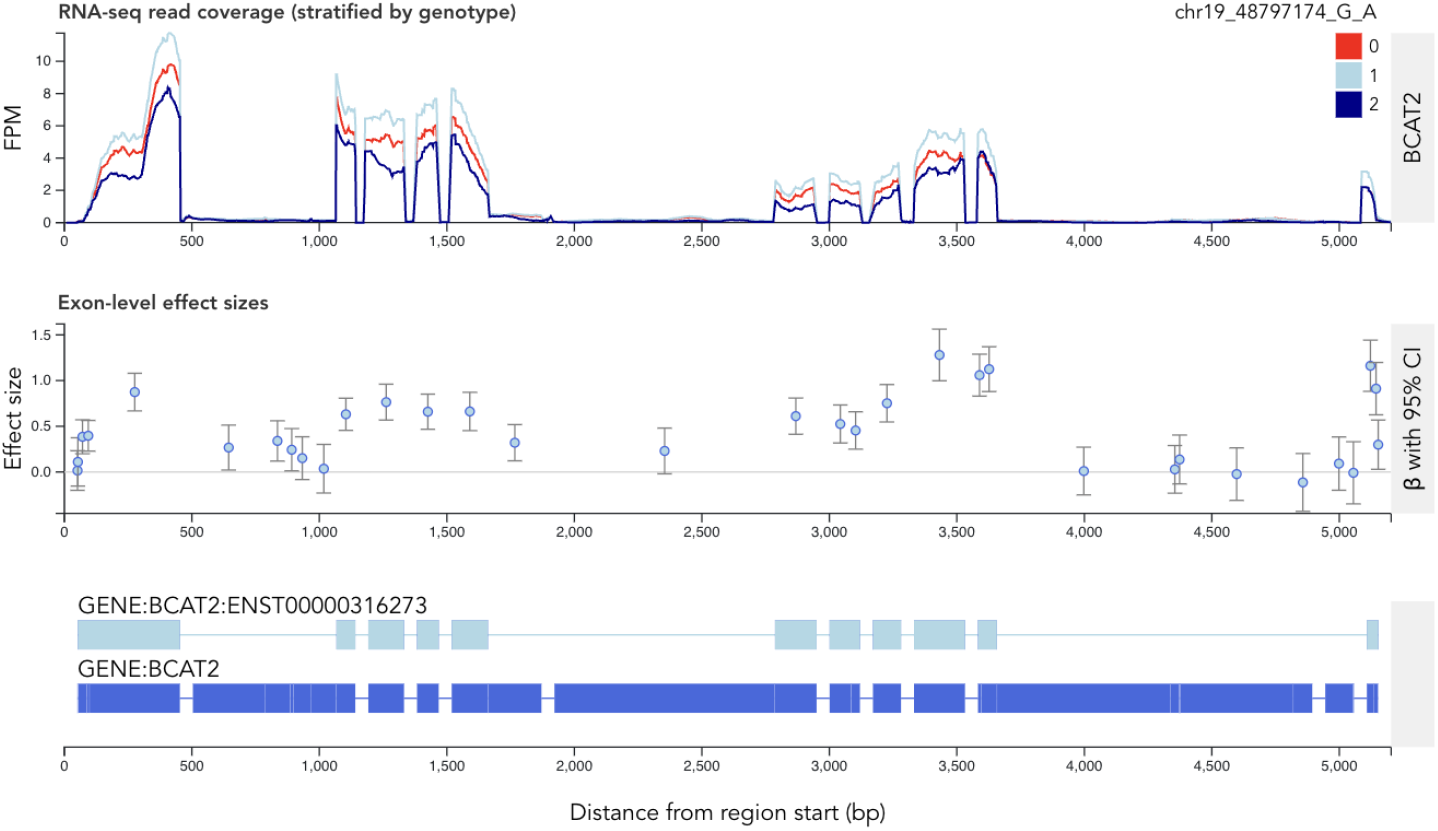
Visualisation of the *BCAT2* expression QTL signal. RNA-seq read coverage across the *BCAT2* gene in the FUSION adipose tissue dataset (QTD000090) stratified by the genotype of the lead eQTL variant (rs35230038). The interactive plot can be viewed in the ELIXIR-Estonia eQTL Catalogue Browser (<u>link</u>).

## Notes

https://github.com/mjesse-github/gpu-coloc

### Competing Interest Statement

The authors have declared no competing interest.

### Summary of Updates

We have edited the Introduction and Results sections of the manuscript to clarify how gpu-coloc relates to existing coloc.abf and coloc.susie implementations.

